# LittleBrain: a gradient-based tool for the topographical interpretation of cerebellar neuroimaging findings

**DOI:** 10.1101/400416

**Authors:** Xavier Guell, Mathias Goncalves, Jakub R Kaczmarzyk, John DE Gabrieli, Jeremy D Schmahmann, Satrajit S Ghosh

## Abstract

Gradient-based approaches to brain function have recently unmasked fundamental properties of brain organization. Diffusion map embedding analysis of resting-state fMRI data revealed a primary-to-transmodal axis of cerebral cortical macroscale functional organization. The same method was recently used to analyze resting-state data within the cerebellum, revealing for the first time a sensorimotor-fugal macroscale organization principle of cerebellar function. Cerebellar gradient 1 extended from motor to non-motor task-unfocused (default-mode network) areas, and cerebellar gradient 2 isolated task-focused processing regions. Here we present a freely available and easily accessible tool that applies this new knowledge to the topographical interpretation of cerebellar neuroimaging findings. LittleBrain generates scatterplots that illustrate the relationship between cerebellar data (e.g., volumetric patient study clusters, task activation maps, etc.) and cerebellar gradients 1 and 2. This novel method of data mapping provides alternative, gradual visualizations that complement discrete parcellation maps of cerebellar functional neuroanatomy. We present application examples to show that LittleBrain can also capture subtle, progressive aspects of cerebellar functional neuroanatomy that would be difficult to visualize using conventional mapping techniques. Download and use instructions can be found at https://xaviergp.github.io/littlebrain.

## INTRODUCTION

Neuroimaging research has greatly improved our knowledge of brain function and structure in health and disease. It has revealed that specific brain regions are associated with specific neurological functions, a territory of knowledge that was previously accessible only through the study of patients with brain injury. It has also provided the first compelling evidence that neuropsychiatric and neurodevelopmental disorders are associated with disruptions in brain structure and function, described brain organization features that can change from subject to subject, and provided new measures that may predict future human behavior such as treatment response in neurology and psychiatry (1).

Topographic interpretation of neuroimaging findings is crucial to the neuroimaging field’s mission to understand the nervous system and alleviate suffering in neurological and psychiatric disorders. Studies often reveal brain activation related to a specific aspect of cognitive processing or brain structural changes associated with a neurological disorder. These findings immediately lead to a topographical interpretation question - what does the distribution of these findings mean? One approach to answer this question is to delineate brain regions and interpret the distribution of these findings based on their overlap with these regions. For example, one may parcellate the cerebral cortex into seven distinct networks (somatomotor, visual, ventral and dorsal attention, frontoparietal, limbic, and default-mode network) (2). The topographical distribution of a cerebral cortical neuroimaging cluster may then be interpreted based on its overlap with these seven regions. Here we propose a different approach to the topographical interpretation of neuroimaging findings based on the utilization of continuous maps (or “gradients”) rather than discrete brain maps.

Several recent studies have adopted gradient-based approaches to describe brain organization. Margulies and colleagues (3) used diffusion map embedding (4) to extract functional gradients from fMRI resting-state data and described two gradients that offer a continuous measure of cerebral cortical functional organization. Gradient 1 extended from primary cortices to transmodal association areas (default-mode network), and gradient 2 extended from primary visual to primary motor/somatosensory and auditory cortical areas (3). A recent review (5) related these findings to other gradient-based analyses of cerebral cortical T1 intensity (a proxy measure of myelin content) (6), neuroimaging measures of semantic representation (7), and neuroimaging measures of variations in the timescale of event representation (8). Diffusion map embedding was recently used to analyze resting-state data within the cerebellum, revealing for the first time a sensorimotor-fugal macroscale organization of cerebellar function (9). Cerebellar gradient 1 extended from motor to non-motor task-unfocused (default-mode) areas, and cerebellar gradient 2 progressed from task-focused to task-unfocused processing regions (9). Gradient-based approaches to brain organization have thus provided novel insights into the functional organization of the brain.

Here we explore the application of this new knowledge to the topographical interpretation of cerebellar neuroimaging findings. Specifically, we aimed to develop a user-friendly tool that ought to facilitate the adoption of a novel gradient-based cerebellar data visualization approach in the neuroimaging community. To achieve this, we minimized installation requirements, maximized user-friendly features, and also empowered experienced users to visualize and modify any features of our tool.

The cerebellum has recently become scientifically relevant not only for the study of motor processes, but also for the study of all complex neurological functions. A large body of literature has shown that the cerebellum is anatomically connected to motor as well as non-motor aspects of the extra-cerebellar structures, that the cerebellum is engaged in numerous motor as well as non-motor aspects of task activation and functional connectivity measures in neuroimaging studies, and that isolated cerebellar injury generates not only a cerebellar motor syndrome but also a cerebellar cognitive affective syndrome (10,11,20–29,12,30,13–19). Similarly, the cerebellum has become clinically relevant for the study of not only primary cases of cerebellar injury or degeneration, but also neurology and psychiatry as a whole. Numerous studies have revealed that many neurological and psychiatric diseases that impair cognitive and affective processing include cerebellar functional and structural abnormalities (31–39). In this way, the present article describes the development of a novel, timely, and relevant neuroscience tool.

## METHODS

### Methods overview

The user of our tool will download and install Docker (https://docker.com). With Docker, the user will download a container that includes a fixed set of all the files and software dependencies required to use our tool. The user will provide, as input, a neuroimaging file that contains cerebellar data. Our tool will then generate a plot that illustrates the relationship between the spatial distribution of these data and the two principal functional gradients of the cerebellum. These two gradients were developed in a previous study by Guell and colleagues (9). Gradient 1 represents a gradual transition from motor to task-unfocused cognitive processing. Gradient 2 isolates task-focused cognitive processing (**Fig. 1**). Cerebellar voxels that are close in the gradient 1 and gradient 2 dimensions have similar functional connectivity patterns. Cerebellar voxels that are distant in the gradient 1 dimension have distinct functional connectivity patterns, specifically as determined by their relationship to the connectivity patterns of motor as opposed to task-unfocused cognitive processing areas of the cerebellum. Cerebellar voxels that are distant in the gradient 2 dimension have distinct functional connectivity patterns, specifically as determined by their relationship to the connectivity patterns of task-focused cognitive processing areas as opposed to the rest of the areas of the cerebellum.

**Figure 1.**
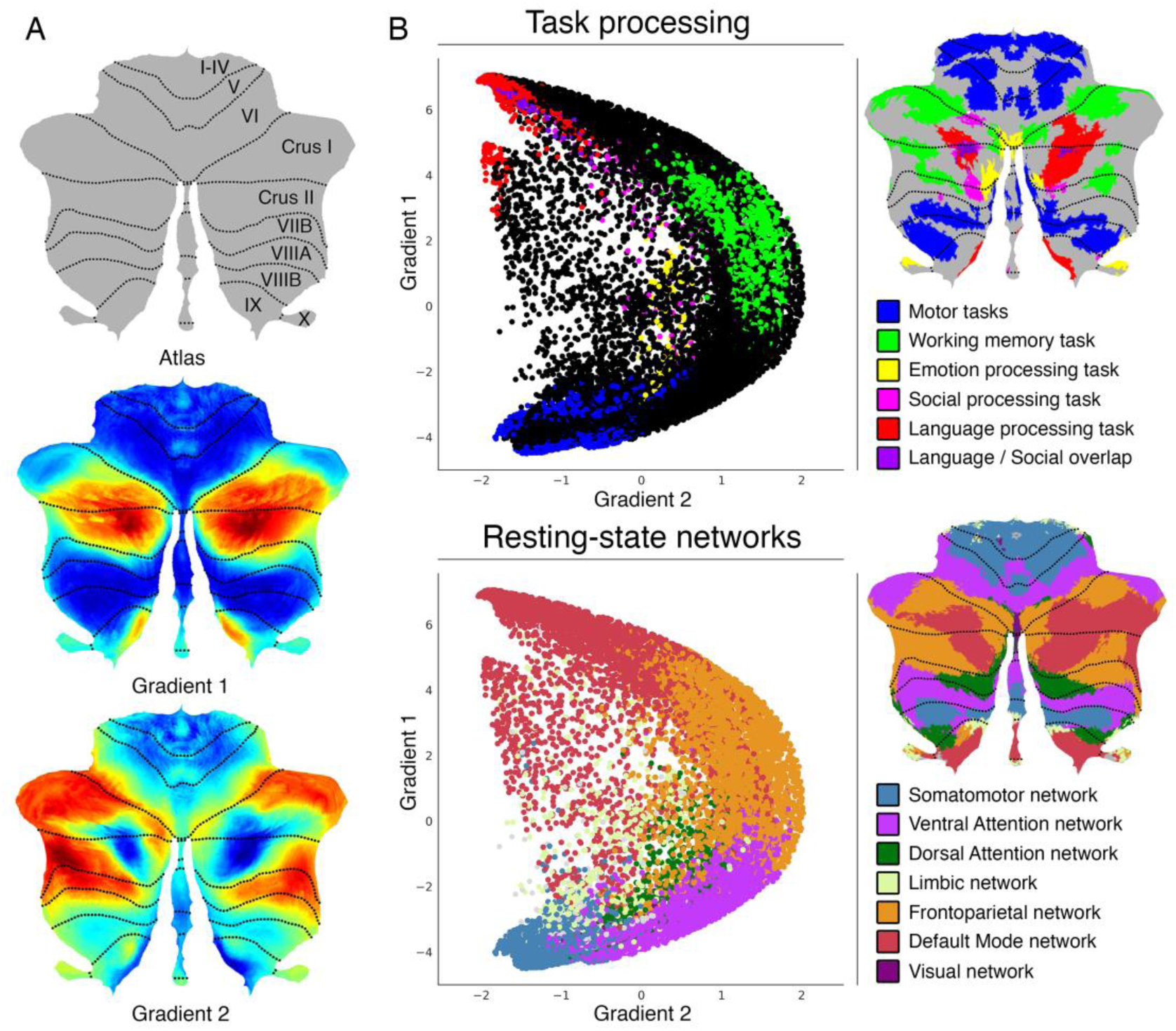
Cerebellum functional gradients and their relationship with discrete task activation and resting-state maps. Adapted with permission from Guell et al., 2018 (9). Gradient 1 extends from motor to task-unfocused cognitive processing regions. Specifically, gradient 1 extends from language task (story listening minus math task contrast, isolating task-unfocused cognitive processing) / Default Mode Network regions to motor regions. Gradient 2 isolates task-focused cognitive processing. Specifically, gradient 2 isolates working memory task processing / task-positive cognitive network areas. **(A)** Gradients 1 and 2 are shown in a colorscale from blue to red. **(B)** A scatterplot of gradient 1 and 2 illustrating their relationship with task (from Guell et al., 2018 (40)) and resting-state maps (from Buckner et al., 2011 (41)). Each dot corresponds to a cerebellar voxel, position of each dot along x and y axis corresponds to position along Gradient 1 and Gradient 2 for that cerebellar voxel, and color of the dot corresponds to task activation (top) or resting-state network (bottom) associated with that particular voxel.

### Input

Neuroimaging data provided by the user must be registered to MNI152 space and be in NIfTI (.nii) format. This NIfTI file may include clusters outside of the cerebellum, and clusters within the cerebellum may extend outside the cerebellum or inside cerebellar white matter - these portions of the data will be removed as part of our preprocessing algorithm. Further, the input does not need to be thresholded or binarized - the user will have the option to select a threshold value when selecting the input, and thresholding and binarization will be performed automatically.

### Data preprocessing

Preprocessing steps include (i) binarization (and thresholding if selected by the user) using FSL’s fslmaths (42), (ii) downsampling to 2mm resolution using FSL’s FLIRT (FMRIB’S Linear Image Registration Tool) (42–44), (iii) adding or deleting slices to match the Human Connectome’s mask files using FSL’s fslroi (42), (iv) masking according to the cerebellum gray matter mask from the Human Connectome Project using Freesurfer’s mrimask (45), so that extra-cerebellar structures as well as white matter aspects of the cerebellum are removed, (v) transforming image size characteristics of the image to match the Human Connectome Project image size characteristics using FSL’s FLIRT (42–44), and (vi) reformatting the image from NIfTI to CIFTI dscalar format using Workbench Command’s -cifti-create-dense-from-template (46).

### Gradients map formation

Following pre-processing, Python-based calculations generate arrays corresponding to gradient 1 and gradient 2 values (as provided by Guell et al., 2018 (9)) for each cerebellar voxel, as well as an equally-sized array consisting of either 1 or 0 values for each cerebellar voxel equivalent to the thresholded and binarized cerebellar data provided by the user. A two-dimensional scatterplot is then generated using matplotlib and seaborn, where each dot in the scatterplot represents a cerebellar voxel, position of each dot along x and y axis corresponds to position along gradient 1 and gradient 2 for that cerebellar voxel, and dots shown in red correspond to the voxels that are included in the cerebellar cluster provided by the user. Before re-starting the process to generate a new gradients map, intermediate files are automatically removed.

### Quality control

While we were unable to identify cases where an MNI-registered NIfTI file was incorrectly transformed, it remains a possibility that preprocessing steps described in this section might result in an erroneous position of the transformed cluster. Extra-cerebellar and cerebellar white matter portions of the data should be removed, but portions of the cluster in cerebellar grey matter should not change their location after preprocessing. The user will have the option to visualize two glass-brain images automatically generated using niwidget to make sure that no erroneous transformations have occurred. Restated, the user will have the option to visualize that position of cerebellar grey matter data has not changed as a result of preprocessing.

### Docker container: simplicity and reproducibility

The processing steps described here require multiple software packages (FSL (42), Freesurfer (45), Connectome Workbench (46), Python 3 and multiple Python packages). The requirement to install these programs would decrease the accessibility of our tool. Further, software updates in these programs might disrupt correct functioning of our tool in the future, or generate replication failures across laboratories. To address these issues, we packaged our tool and all its files and software dependencies in a “Docker container” (47) using neurodocker (48). Docker (47) is an open source project that allows the encapsulation of multiple software packages and files into a single “container”. This container can be stored online in a public repository, be easily downloaded by users worldwide, and guarantees that any computer will be able to perform all operations inside the container in the same way. Users will need to install Docker on their computers, and will be able to use our tool by simply downloading and running one single Docker container. No software paid subscriptions are required to use our tool. However, if you are considering commercial use of FSL, please consult their license at https://fsl.fmrib.ox.ac.uk/fsl/fslwiki/Licence. Our Docker container has been minimized using reprozip (49), and hence includes only those aspects of FSL, Freesurfer, and Connectome Workbench that are necessary to perform the functions of LittleBrain. This reduced the size of LittleBrain to 2.5 gigabytes (compared to 12 gigabytes if we had not performed this minimizing step).

### Jupyter Notebook: transparency and flexibility

Jupyter Notebook (50) is a browser application that allows users to code interactively, display graphics, and enter user input. In this way, while our tool displays figures (gradient 1/2 scatterplot, quality control output), accepts user input using text boxes, and provides an intuitive interface, it also exposes the code for each step of the process, and allows users with coding experience to visualize and modify any feature of our tool. Examples of the flexibility that Jupyter Notebook provides are given in the following section.

## RESULTS

In this section we describe the architecture and features of LittleBrain (**Fig. 2**) and present two example analyses. Three additional examples are then shown to illustrate alternative visualization strategies that experienced users might implement by using the interactive Jupyter notebook.

**Figure 2.**
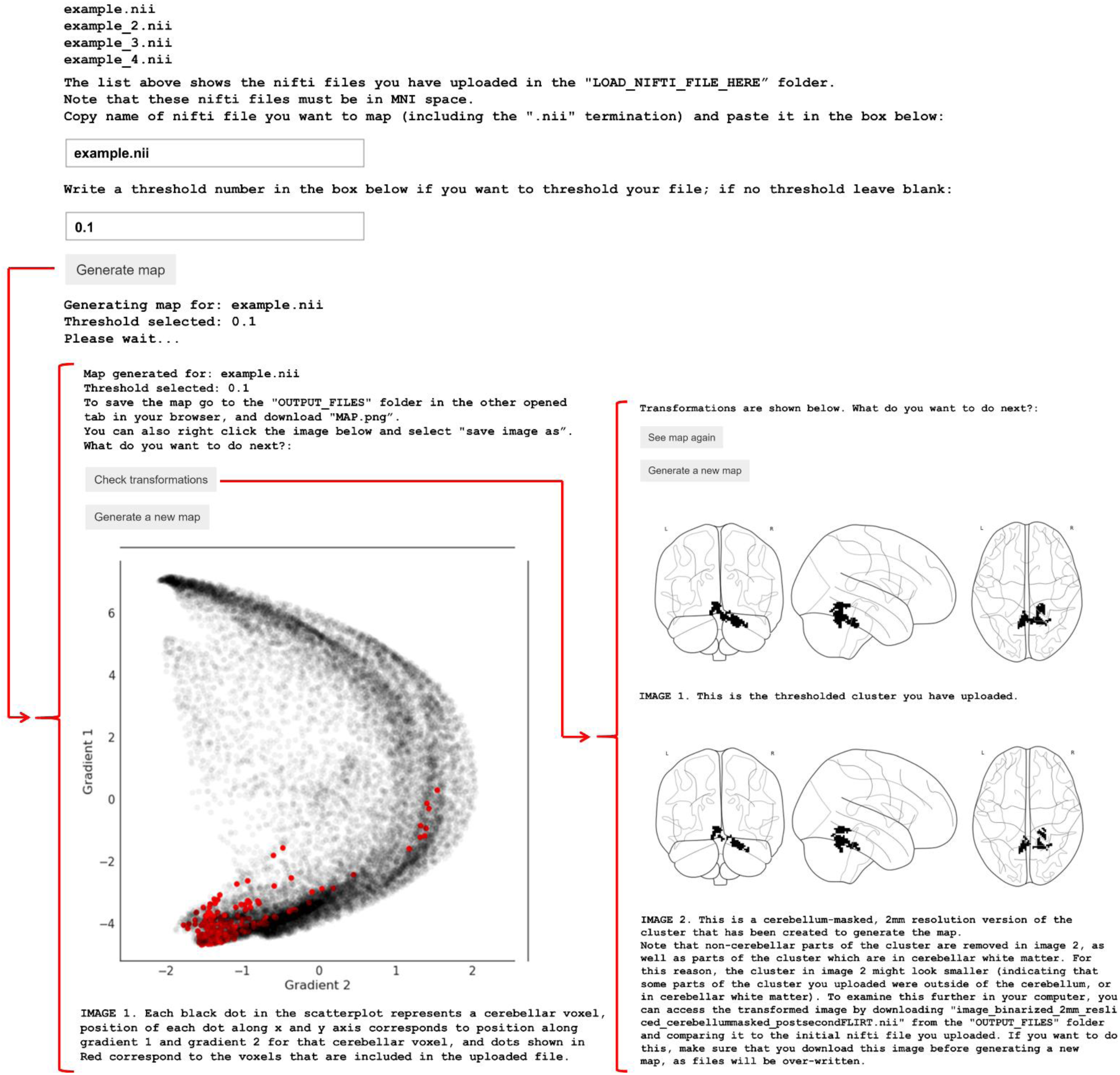
Visualization of the user interface of LittleBrain. After uploading NIfTI files containing cerebellar data and running the Jupyter notebook, LittleBrain shows a list of all inputs that have been uploaded. The user enters the name of the input NIfTI file that will be mapped and may also enter a threshold value to be applied to the NIfTI file. After clicking “Generate gradients map” and waiting a few seconds, the screen will display a scatterplot illustrating the relationship of cerebellar data with gradients 1 and 2 from Guell et al., 2018 (9). In the scatterplot, each dot corresponds to a cerebellar voxel, position of each dot along x and y axis corresponds to position along Gradient 1 and Gradient 2 for that cerebellar voxel, and red dots indicate which cerebellar voxels are included in the data provided by the user after thresholding and binarization. The user will then have the option to quality control by clicking “Check transformations”. This option displays two glass-brain images corresponding to the binarized and thresholded cluster before (top image) and after (bottom image) preprocessing transformations have occurred. Extra-cerebellar and cerebellar white matter portions of the data should be removed, but portions of the cluster in cerebellar grey matter should not change their location after preprocessing.

### Application example 1

An unpublished study from our laboratory used multi-voxel pattern analysis (MVPA) to identify brain regions with abnormal whole-brain functional connectivity patterns in a patient with a retrocerebellar arachnoid cyst compressing the cerebellum and a life-long history of psychiatric and neurodevelopmental symptoms. The patient’s data was compared to 36 healthy controls. This analysis revealed a cluster in cerebellar lobules I through VI (**Fig. 3A**). Lobules I-VI are engaged in motor processing. Multiple studies also indicate a non-motor role of some aspects of lobule VI (21,24,25,40). A cerebellar resting-state network map (41) revealed inclusion of somatosensory network as well as some aspects of ventral attention network (**Fig. 3B**), indicating that this cluster may include predominantly motor (somatomotor network) but also non-motor task-focused cognitive processing regions (ventral attention network). LittleBrain provided an unparalleled, clear visualization of this concept (**Fig. 3C**). Most voxels of the cerebellar cluster had low gradient 1 / low gradient 2 values, indicating a strong inclusion of motor processing regions. As expected, a few voxels extended towards task-focused cognitive processing regions (i.e. high gradient 2 values). LittleBrain can thus provide alternative, gradual visualizations that complement discrete parcellation maps of cerebellar functional neuroanatomy. This example can be reproduced by mapping the “example.nii” file included in LittleBrain.

**Figure 3.**
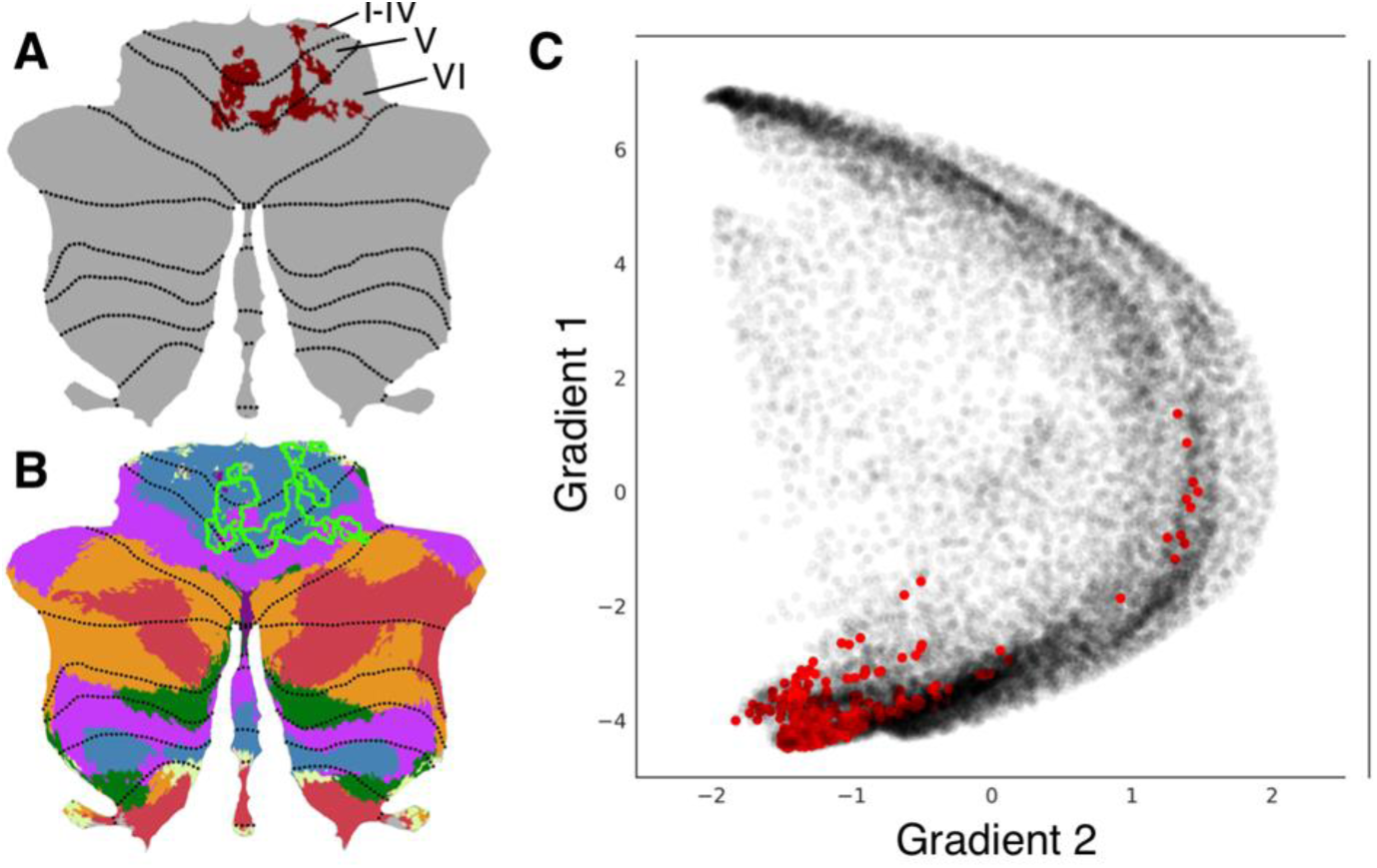
LittleBrain provides alternative, gradual visualizations that complement discrete parcellation maps of cerebellar functional neuroanatomy. **(A)** Cerebellar MVPA cluster (red) was located in lobules I-VI. **(B)** Overlap of this cluster (green outline) with a resting-state networks map (41) revealed inclusion of somatosensory network (blue) as well as some aspects of ventral attention network (purple). **(C)** Each dot corresponds to a cerebellar voxel, position of each dot along x and y axis corresponds to position along gradients 1 and 2 for that cerebellar voxel, and red dots correspond to those cerebellar voxels that are included in the cerebellar cluster shown in A. This provides an unparalleled, clear visualization of the concept that this cerebellar cluster is mainly localized in motor processing regions (low gradient 1 / low gradient 2 values) but also encroaches task-positive cognitive processing areas (high gradient 2 values).

### Application example 2

A study by Lesage and colleagues (51) performed two localizer task experiments in a group of 18 healthy participants. The *“Semantic” working memory task* contrasted a 1-back condition (while watching pictures of objects, press a button if the semantic category of the present image is the same as the semantic category of the previous image; e.g., press the button if the picture of a sailing boat is followed by a picture of a motor boat, but not if a picture of a sailing boat is followed by a picture of a fish) minus a 0-back condition (similar pictures are shown, and participants are required to press the button when a target image appears, e.g. press the button every time the picture of a sailing boat appears in the screen). Of note, subjects were previously familiarized with 10 stimulus categories: cycles, birds, boats, dogs, fish, fruits, buildings, shoes, tools, and furniture. The *“Punjabi” working memory task* contrasted a 1-back condition (while watching pictures of incomprehensible Punjabi alphabet pseudowords, press a button if the pseudoword of the present image is the same as the pseudoword of the previous image) minus a 0-back condition (similar pictures are shown, and participants are required to press the button when a target pseudoword appears) (see **Fig. 4**). Task contrast cluster-extent correction based thresholded maps from these analyses were downloaded from Neurovault (52), and we performed additional thresholding (values>6) to better visualize differences between the two maps.

**Figure 4.**
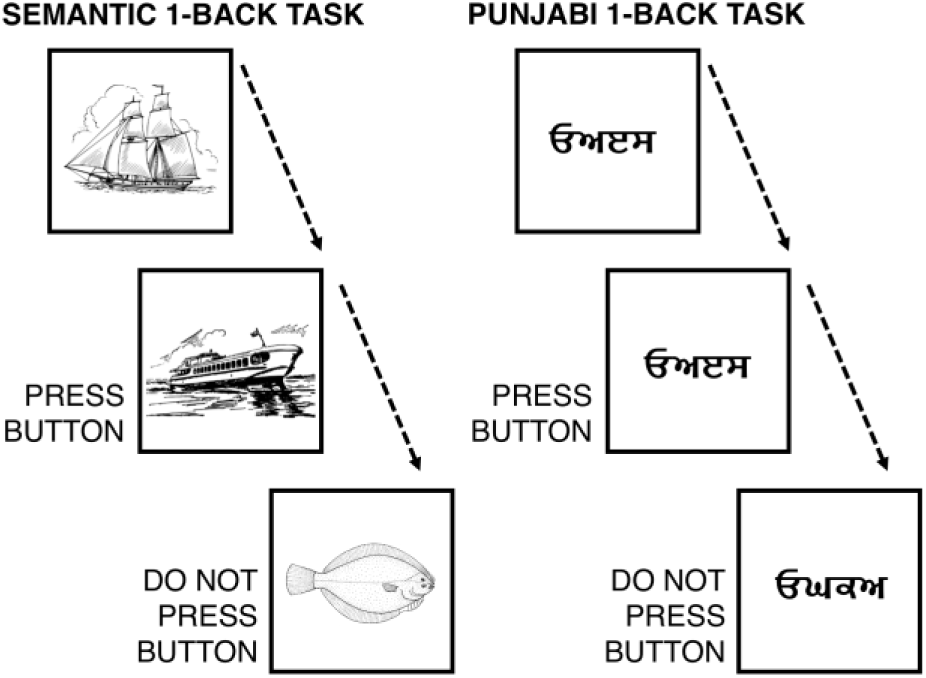
Scheme of semantic 1-back condition and visual (punjabi) 1-back condition. Both tasks engaged non-motor aspects of the cerebellum (53), specifically lobules VI, Crus I/II, and VIIB (**Fig. 5**). A cerebellar resting-state networks map (41) revealed engagement of ventral/dorsal attention and frontoparietal networks in both tasks, as well as marginal encroachment of default mode network areas in the Semantic task only (**Fig. 5**). Approximation to default mode network regions in the semantic task might reflect partial engagement of unfocused cognitive processing regions; perhaps contrasting with higher focused cognitive demands in the Punjabi task, which was located in aspects of task positive networks (dorsal/ventral attention, frontoparietal) that were slightly more distant from cerebellar default-mode network areas. LittleBrain offered an unparalleled visualization of this concept (**Fig. 5**). Punjabi task was located at high gradient 2 areas (corresponding to focused cognitive processing). Semantic task was located at similar regions but extending towards higher gradient 1 values (i.e., more unfocused cognitive processing regions) and lower gradient 2 values (i.e., less focused cognitive processing regions). In this way, Semantic task may engage cerebellar regions which are slightly less involved in focused cognitive processing when compared to Punjabi task, and slightly more involved in unfocused cognitive processing. LittleBrain can thus capture subtle, progressive aspects of cerebellar functional neuroanatomy that would be difficult to visualize using conventional discrete parcellations of cerebellar functional neuroanatomy. Of note, comparison between gradients maps may be supplemented by additional statistical analyses – this possibility is explored in the following section.

**Figure 5.**
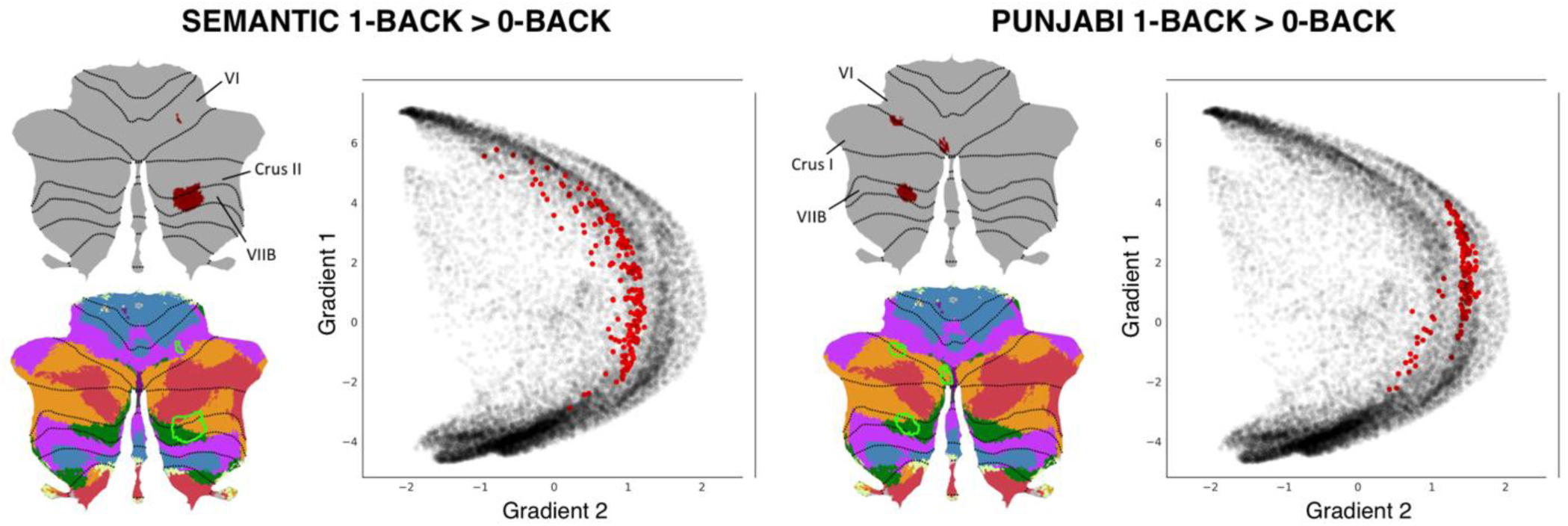
LittleBrain can capture subtle, progressive aspects of cerebellar functional neuroanatomy that would be difficult to visualize using conventional discrete parcellations of cerebellar functional neuroanatomy. Both tasks engaged lobules VI, Crus I/II, and VIIB. Both tasks engaged ventral/dorsal attention (green and purple) and frontoparietal (orange) networks (i.e., focused cognitive processing areas). Marginal aspects of default mode network (red) (i.e., unfocused cognitive processing areas) were included in Semantic task only. Semantic task may engage cerebellar regions which are slightly less involved in focused cognitive processing when compared to Punjabi task, and slightly more involved in unfocused cognitive processing. LittleBrain provides an unparalleled visualization of this concept – Semantic task reveals higher gradient 1 values (i.e., more unfocused cognitive processing regions) and lower gradient 2 values (i.e., less focused cognitive processing regions).

### Flexibility examples

Our public source code repository (https://github.com/xaviergp/littlebrain) includes several examples of alternative visualization and analysis strategies that users with coding experience might implement by interacting with the Jupyter notebook. These include (i) mapping multiple cerebellar clusters at once, each with a different color (**Fig. 6A**), (ii) extracting gradient 1 and 2 values for each voxel included within a cerebellar cluster and performing statistical analyses using these data (**Fig. 6B**), and (iii) mapping cerebellar clusters using gradients other than 1 and 2 from Guell et al., 2018 (9) (e.g. using gradients 1 and 3) (**Fig. 6C**). LittleBrain is therefore easy to install and use, but also allows the experienced user to develop complex novel visualization or analysis strategies. These alternative visualizations and analyses can be performed using Python packages included in our LittleBrain Docker container such as seaborn and scipy. Because LittleBrain is hosted publicly, users who develop new strategies may push their contributions back to the LittleBrain source code, or to the examples in the LittleBrain repository.

**Figure 6.**
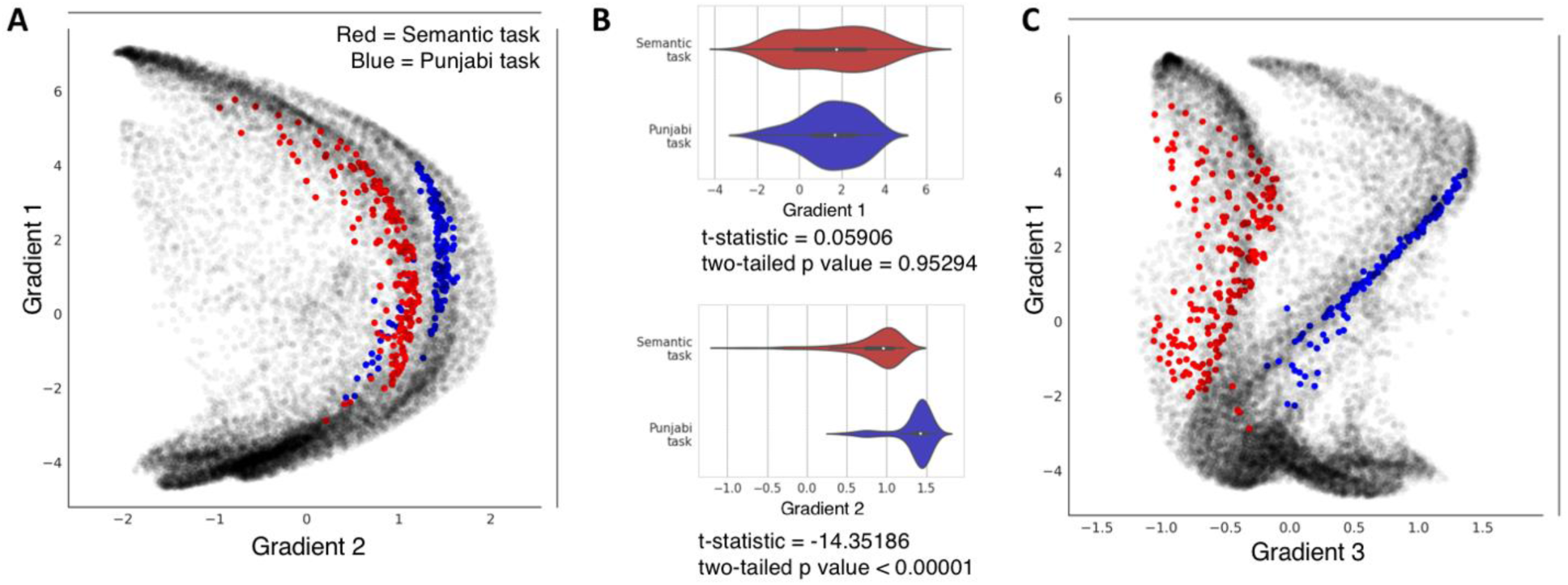
Flexibility examples of LittleBrain using data from our previous example (Fig. 5). **(A)** Plotting two task maps in a single gradients map allows a better visualization of the relationship between them. Semantic task cluster is shown in red, Punjabi task cluster in shown in blue. **(B)** It is possible to extract gradient 1 and gradient 2 values from each map, generate plots using these data (such as violin plots, as shown here), and also perform statistical tests using these data (such as t-tests, as shown here). This allowed us to detect, for instance, that gradient 2 but not gradient 1 differences between the two maps were statistically significant. **(C)** It is possible to use gradients other than gradients 1 or 2. For example, gradient 3 represents left-right hemisphere asymmetries in nonmotor cerebellar areas (see discussion in Guell et al., 2018 (9) regarding gradient 3). Accordingly, gradient 3 separates our two task maps given that they both engage non-motor processing areas, but are located at opposite cerebellar hemispheres (see Fig. 5; Semantic task activation map is located in right cerebellar hemisphere while Punjabi task activation map is located in left cerebellar hemisphere).

### LittleBrain Manual

A manual with detailed installation and use instructions is available at https://xaviergp.github.io/littlebrain.

## DISCUSSION

We present LittleBrain, a novel neuroimaging data visualization tool that offers an unparalleled perspective to the topographical interpretation of cerebellar neuroimaging findings. Gradient-based approaches to brain function have recently unmasked fundamental properties of brain organization. LittleBrain applies this new knowledge to the topographical interpretation of cerebellar neuroimaging findings. It is an easily accessible, user-friendly tool that can also be interactively modified by experienced users.

Using real data examples, we have shown that LittleBrain can provide alternative, gradual visualizations that complement traditional discrete parcellation maps of cerebellar functional neuroanatomy. We have also shown that LittleBrain can capture subtle, progressive aspects of cerebellar organization that would be difficult to visualize using conventional discrete parcellations. LittleBrain may thus improve the topographical interpretation of cerebellar neuroimaging findings, in a context where the cerebellum is receiving increased attention as a necessary component for understanding virtually all complex brain functions in health and disease.

Future research might expand the applications of the present work. For example, similar tools could be developed for the analysis of data from other brain regions. Margulies and colleagues (3) described functional gradients in the cerebral cortex – LittleBrain could be extended to operate on cerebral cortical data. Similarly, future tools may use gradient maps developed using methods other than resting-state diffusion map embedding, such as continuous semantic maps as developed by Huth and colleagues (7), continuous event timescale representation maps as developed by Baldassano and colleagues (8), or even continuous maps related to structural rather than functional brain properties (6).

Novel methods of analysis might also emerge from this new method of data visualization. As shown in **Fig. 6B**, statistical analyses might be performed using gradient values extracted from cerebellar clusters. Functional neuroanatomical differences between cerebellar clusters may therefore not only be visualized, but also quantified. Future research might explore the utility of this approach and determine the optimal strategy for such analyses.

Taken together, LittleBrain is a novel and fully functional tool that may significantly improve cerebellar neuroimaging data interpretation. It is accessible to the novice user, and transparent and flexible to the experienced user. It links novel developments in gradient-based functional neuroanatomy to the topographical interpretation of cerebellar neuroimaging findings, unmasking otherwise inaccessible functional properties of cerebellar data topography. It resonates with a paradigm shift in the appreciation of the relevance of the cerebellum in neuroscience, and with an increased attention to gradient-based approaches to brain function in the neuroimaging community.

## ACKNOWLEDGEMENTS

This study was supported in part by La Caixa Banking Foundation (XG), the MGH ECOR Fund for Medical Discovery Postdoctoral Fellowship Award (XG), the MINDLink foundation (JDS), and National Institutes of Health R01 EB020740 (SSG) and P41 EB019936 (SSG). The funders had no role in study design, data collection and analysis, decision to publish, or preparation of the manuscript. The authors want to thank Paul Wighton for assistance when building Docker images, and Theodore LaGrow for participating in our toolbox name brainstorming.

